# Multiple resource limitations explain biomass-precipitation relationships in grasslands

**DOI:** 10.1101/2021.03.09.434527

**Authors:** Siddharth Bharath, Peter B. Adler, Philip A. Fay, Eric W. Seabloom, Yann Hautier, Lori Biederman, Miguel N. Bugalho, Maria Caldeira, Anu Eskelinen, Johannes M.H. Knops, Rebecca McCulley, John Morgan, Sally A Power, Anita C. Risch, Martin Schuetz, Carly J. Stevens, Timothy Ohlert, Risto Virtanen, Elizabeth T. Borer

## Abstract

Interannual variability in grassland primary production is strongly driven by precipitation, nutrient availability and herbivory, but there is no general consensus on the mechanisms linking these variables. If grassland biomass is limited by the single most limiting resource at a given time, then we expect that nutrient addition will not affect biomass production at arid sites. We conducted a distributed experiment manipulating nutrients and herbivores at 44 grassland sites in 8 regions around the world, spanning a broad range in aridity. We estimated the effects of 5-11 years of nutrient addition and herbivore exclusion treatments on precipitation sensitivity of biomass (proportional change in biomass relative to proportional change in rainfall among years), and the biomass in the driest year (to measure treatment effects when water was most limiting) at each site. Grazer exclusion did not interact with nutrients to influence driest year biomass or sensitivity. Nutrient addition increased driest year biomass by 74% and sensitivity by 0.12 (proportional units), and that effect did not change across the range of aridity spanned by our sites. Grazer exclusion did not interact with nutrients to influence sensitivity or driest year biomass. At almost half of our sites, the previous year's rainfall explained as much variation in biomass as current year precipitation. Overall, our distributed fertilization experiment detected co-limitation between nutrients and water governing grasslands, with biomass sensitivity to precipitation being limited by nutrient availability irrespective of site aridity and herbivory. Our findings refute the classical ideas that grassland plant performance is limited by the single most limiting resource at a site. This suggests that nutrient eutrophication will destabilize grassland ecosystems through increased sensitivity to precipitation variation.

## 1 Introduction

The productivity of grassland ecosystems around the world is strongly driven by precipitation. Anthropogenic global change is expected to increase interannual variation in precipitation (Fischer et al., 2013). Aboveground plant biomass in grasslands (henceforth “biomass”) is additionally governed by soil resource availability (Fay et al., 2015) and consumption by grazers (Borer, Seabloom, et al., 2014), and both of these should influence how variation in precipitation over time affects biomass production at a site (Huxman et al., 2004; Irisarri et al., 2016). Understanding the nature of this temporal relationship across different sites and regions of the world is crucial for predicting how rainfall variability will interact with other global changes such as nutrient eutrophication (Stevens et al., 2015).

The temporal relationship between annual aboveground biomass and annual precipitation in grasslands has been extensively measured in long-term ecological studies (Hsu et al., 2012; Sala, Gherardi, et al., 2012; Sala, Parton, et al., 1988). Two aspects of this relationship at a site are important to this study. Firstly, the 'sensitivity’ of biomass to interannual precipitation variation at a site measures how much biomass changes for a given change in rainfall among years. Secondly, the biomass produced during the driest year for a specific site, and the effects of resource addition treatments on that, gives an understanding of the importance of other resources when water is most limiting (see Figure 1a). Both of these measures (sensitivity and driest year biomass) are expected to vary among sites, especially with relation to site aridity (Bai et al., 2008; Hsu et al., 2012; Huxman et al., 2004). Since limitation by water should be stronger in arid sites, the sensitivity of biomass to precipitation should decrease as we move from arid to mesic sites (Huxman et al., 2004; Sala, Gherardi, et al., 2012, Figure 1b). Mean grassland biomass and driest year biomass should increase as we move from arid to mesic sites (Sala, Gherardi, et al., 2012; Sala, Parton, et al., 1988), though there can be variations in these patterns among grassland regions (Bharath et al., 2020; O’Halloran et al., 2013).

However, precipitation-production patterns may depend on nutrient limitation. Most grasslands are not only limited by water, but also by the supply of available soil nutrients; e.g., nitrogen or phosphorus (Fay et al., 2015). The degree of nutrient limitation is expected to vary based on the aridity of the site (Yahdjian et al., 2011). Experimental nutrient addition should alleviate limitations, thus increase the sensitivity of biomass to precipitation in mesic, but not arid sites (Figure 1, Bharath et al., 2020; Huxman et al., 2004; Wang et al., 2017). Water availability should primarily limit biomass in dry years, and therefore nutrient addition should have little effect on driest year biomass at arid sites, but increase biomass at mesic sites (Yahdjian et al., 2011). Alternatively, if biomass production of a plant community is equally constrained by multiple resources (Rastetter and Shaver, 1992), nutrient addition can increase biomass production even in dry years at arid sites (Hooper and Johnson, 1999). Examining how alleviation of nutrient limitation interacts with moisture limitation across time and space enables us to evaluate the utility of the multiple resource limitation framework for understanding grassland productivity.

**Figure 1:**
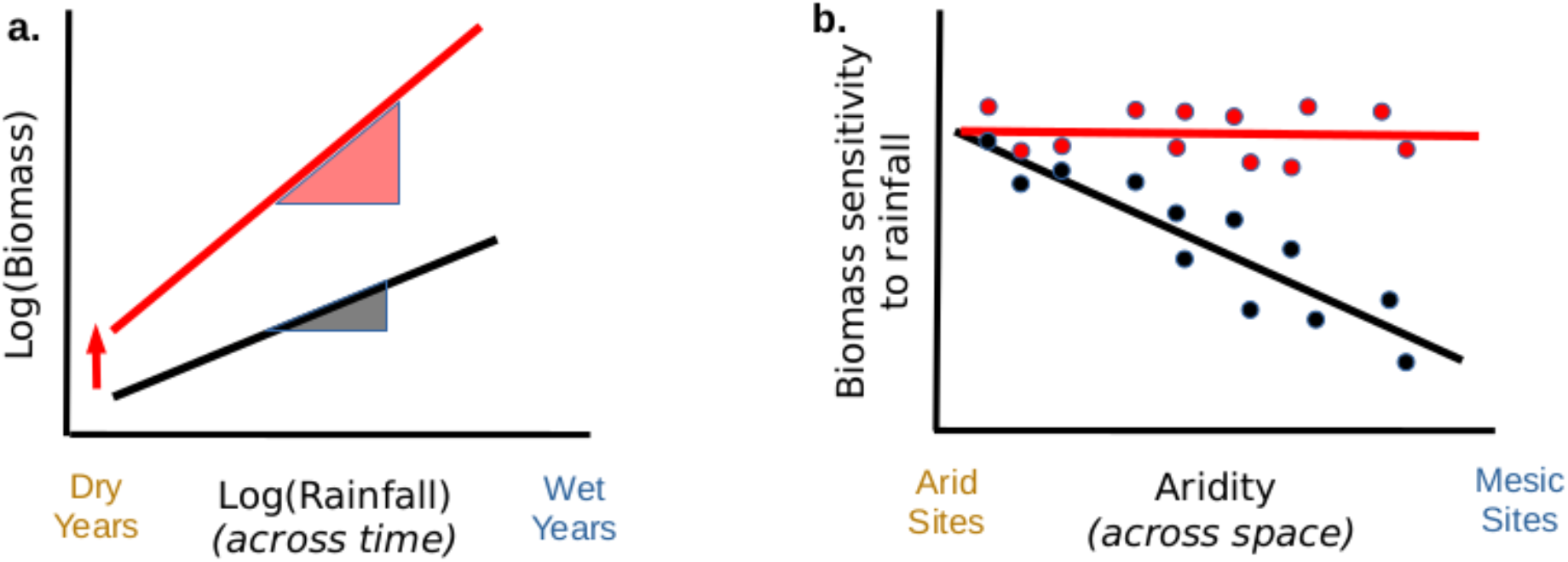
Expectations of single resource limitation driving grassland sensitivity to rainfall, from Huxman et al. (2004) **a.** Nutrient addition is expected to increase both biomass measured in the driest year (bd), and the precipitation sensitivity S (proportional response of biomass to a change in rainfall), estimated from the graph of biomass vs. precipitation across time at a site. Black denotes the relationship in control plots, red in nutrient added plots. **b.** Across space, precipitation sensitivity is expected to decline from arid to mesic sites. Since sensitivity is already maximum at arid sites, nutrient addition is expected to have no effect on sensitivity at arid sites, but a strong effect at mesic sites.

The effects of herbivory on plant biomass should interact with nutrient availability and precipitation in grasslands (Anderson et al., 2018; Frank et al., 2018; McNaughton et al., 1989). The net effect of grazers on biomass across years at a site can alter biomass-precipitation relationships in different ways. Depending on how consumption by grazers changes between wet and dry years, their exclusion might either increase or decrease sensitivity of biomass to precipitation. If grazers at a site are consuming the additional biomass generated by fertilization, their exclusion will increase the effects of nutrients on sensitivity (Gruner et al., 2008). If grazing enhances production by releasing the plant community from light limitation (Borer, Seabloom, et al., 2014), nutrients will have smaller effects on sensitivity when grazers are excluded. In spite of all these possible mechanisms, the generality and degree to which grazing by mammalian herbivores modulates the relationship between grassland biomass and interannual variation in precipitation has not been quantitatively evaluated (Campbell and Stafford Smith, 2000; Frank et al., 2018).

We quantified how precipitation sensitivity and driest year biomass were affected by 5-11 years of continuous nutrient addition in 44 grassland sites around the world. We also tested whether effects of nutrients were altered by the gradient of aridity among sites and the simultaneous experimental exclusion of vertebrate herbivores (at 36 sites). We specifically examined the following predictions derived from the hypothesis that water is among multiple resources that co-limit grassland productivity –

1. Precipitation sensitivity of biomass will decline from arid to mesic sites (Huxman et al., 2004; Sala, Gherardi, et al., 2012).
2. Nutrient addition will increase precipitation sensitivity at mesic sites but not at arid sites (Huxman et al., 2004).
3. Driest year biomass will increase from arid to mesic sites, as the amount of water received in the driest year is higher at mesic sites as compared to arid sites.
4. Nutrient addition will have no effect on driest year biomass at arid sites, yet will have larger effects at mesic sites (Yahdjian et al., 2011).
5. Herbivore exclusion could increase or decrease the effects of nutrients on precipitation sensitivity.

## 2 Methods

### 2.1 Experimental setup

We used data generated within the Nutrient Network (NutNet) experiment, a distributed research cooperative focused on the study of the diversity, productivity, and composition of grasslands worldwide (Borer, Harpole, et al., 2014). Within the network, we selected all sites that had at least 5 years of biomass data from nutrient treatments, and reliable weather data (described below), which came to a total of 44 sites. Nutrient treatment (N, P, K, and micronutrients added) and fencing were crossed in a factorial design to test for the effects of multiple nutrient limitation and grazing on plant composition and ecosystem function. Experimental plots were 5m × 5m in size, with one set of all treatments arranged in a spatial block, and there were 3-6 blocks per site. Nutrient addition rates and sources were: 10 g N m^−2^year^−1^ as timed-release urea [(NH_2_)_2_CO], 10 g P m^−2^year^−1^ as triple-super phosphate [Ca(H_2_PO_4_)_2_], 10 g K m^−2^year^−1^ as potassium sulfate [K_2_SO_4_] and 100 g m^−2^ of a micronutrient mix of Fe (15%), S (14%), Mg (1.5%), Mn (2.5%), Cu (1%), Zn (1%), B (0.2%), and Mo (0.05%). N, P, and K were applied annually; micronutrients were applied once at the start of the experiment to avoid toxicity (see Borer, Harpole, et al., 2014, for details). The goal of nutrient addition was to overcome resource limitation for plant growth. Each plot was sampled annually for aboveground biomass, clipped from two 0.1 m^2^ quadrats per plot, dried to constant mass at 60°C and weighed to the nearest 0.01g.

At 36 of the 44 sites, we established fences designed to exclude aboveground mammalian herbivores larger than 50 g around two plots in each block, one receiving experimental nutrient addition and one used as an ambient nutrient control plot. Fences were 230 cm tall with the lower 90 cm surrounded by 1-cm woven wire mesh. An additional 30-cm outward-facing flange was stapled to the ground to exclude digging animals (such as rabbits or echidnas), although not fully subterranean animals (such as gophers or mole rats). Four strands of barbless wire were strung at equal vertical distances above the wire mesh. Although most sites built fences exactly to these specifications, 5 sites made minor modifications (described in Appendix S2).

### 2.2 Site selection and weather data

We obtained weather data (total precipitation, average monthly maximum and minimum temperatures) from the nearest reliable source, validated by PIs to be representative of weather at their site (detailed in Appendix S2). Weather data started at least 3 years prior to the first biomass observations. For 43 sites, this was obtained as daily resolution data from weather stations, for 1 site this was as monthly resolution globally gridded data (see list of sources and methodology in Appendix S2). We used a modified form of the Hargreaves equation (Droogers and Allen, 2002) to estimate total potential evapotranspiration (PET) at the monthly scale for each site. Water availability metrics over the growing season are better predictors of annual biomass than annual precipitation (Robinson et al., 2013). We summed precipitation over the growing season months at each site (henceforth abbreviated to GSP); the months that constitute the growing season were determined by site PIs (See Appendix S2: Table S2). We defined aridity of a site as the log_2_ ratio of mean GSP divided by mean growing season PET at a site.

### 2.3 Measuring biomass-precipitation relationships

Temporal biomass-precipitation relationships were found to be nonlinear or saturating in many long term studies (Hsu et al., 2012; Rudgers et al., 2018). We examined the relationships between annual aboveground biomass and GSP at each site by log_2_ transforming both variables and then fitting linear models to the data. This allowed us to fit nonlinear relationships, meet assumptions of normality for linear models, and prevent model predictions of negative biomass in dry years.

We fit linear mixed effects models of the following form at each site:

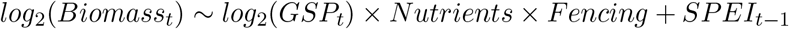

Peak biomass at year *t* (log_2_ transformed) was the response variable. Predictors were GSP, fencing, nutrient addition, as well as all interactions between the these three, allowing both slope and intercept to vary for each treatment at a site. Legacies of the previous year's precipitation can influence biomass by changing the soil seed bank, bud bank, tiller density, soil nutrient pools or soil moisture availability in the current year (Reichmann et al., 2013; Sala, Gherardi, et al., 2012). Therefore, we included the Standardized Precipitation Evapotranspiration Index (SPEI) calculated over the growing season for the previous year as a predictor variable. The SPEI is a normalized metric of water availability in a given year relative to the precipitation and temperature history of the site. This metric is positive if the previous year was wetter than the mean, and negative if it was drier than the mean. We also included a random effect for blocks within sites, to correctly account for the design of our experiment. All analyses were performed in R version 4.0.0 (R Core Team, 2020).

We measured precipitation sensitivity (*S*) as the slope of the relationship between log_2_(Biomass) and log_2_(GSP). A slope value of 1 means that biomass value doubles when precipitation doubles. *S* < 1 indicates that a change in precipitation results in a less than proportional change in biomass, and *S* > 1 indicates a greater than proportional change. Fitted models and parameters are shown in Appendix S1.

Driest year biomass (*b_d_*) was directly estimated as the measured biomass in the driest year during our experiment. We calculated the effects of treatments on this biomass as the log_2_ ratio of biomass in treatment plots over control plots in each block, during that driest year.

We also fit linear relationships to the data of biomass and precipitation (Huxman et al., 2004; Irisarri et al., 2016; Verón et al., 2005), to match earlier studies in the literature. These are reported in Appendix S3.

### 2.4 Examining variation in responses among sites

We then examined the effects of nutrient addition treatments (inside and outside fences) on sensitivity (*S*) and driest year biomass (*b_d_*) along the gradient of aridity (GSP/PET) among our sites. We tested 3 possible models of the relationship between response parameters (y) and GSP/PET (x) –

1. A simple linear regression.
2. A linear regression with different intercepts for each region.
3. A linear regression with different slopes and intercepts for each region.

Since there are differences in uncertainty of the response parameters among sites, we carried out weighted linear regressions where the contribution of a data point to the sums of squares during regression was weighted by the inverse of the standard error of that data point.

Four regions in our study had more than 5 sites each, and were amenable to examination of regional relationships. Thus we first fitted all 3 models to the data of 36 sites, located in the regions of Europe (9 sites), Australia (8 sites), North America Pacific Coast (7 sites) and North America Central Plains (12 sites). This excluded sites in North America Montane West (4 sites), South America (2 sites), sub-Saharan Africa (1 site) and Asia (1 site). For each response variable, we used AIC_c_ based model selection to identify which of these models best describes the data, and report those results. If model selection showed that region was unimportant (models 2 and 3 did not perform well), we then re-fit and reported the results of model 1 on the whole dataset of 44 sites.

We chose this two-stage analyses instead of fitting a linear mixed effect model (MEM) to the global data with random effects for sites. We are interested in examining the variation in biomass-precipitation relationships at many sites. Shrinkage of random parameter estimates towards global means in MEM results in very poor (and in many cases wrong) estimations of biomass-precipitation relationships at individual sites. Global MEM additionally faced convergence issues.

## 3 Results

### Average results across all sites

Across our 44 sites, we found much variation in the shape of biomass-precipitation relationships (Figure 2a). The log-log models used in our study better predicted biomass-precipitation relationships, resulting in 32% less residual variance than the linear models (variance calculated on arithmetic scale for both). The sensitivity of biomass to interannual precipitation at each site (*S*) varied in value from −1.9 to 4.5, with a median value of 0.35. Positive sensitivity values indicate a constant proportional increase in biomass with increasing precipitation. Fifteen out of 44 sites had negative values of sensitivity, indicating that water was not limiting biomass at those sites, or excess water negatively affected biomass. The driest year biomass (*b_d_*) estimated at each site varied from 18 g m^−2^ to 1003 g m^−2^ (Figure 2b).

**Figure 2:**
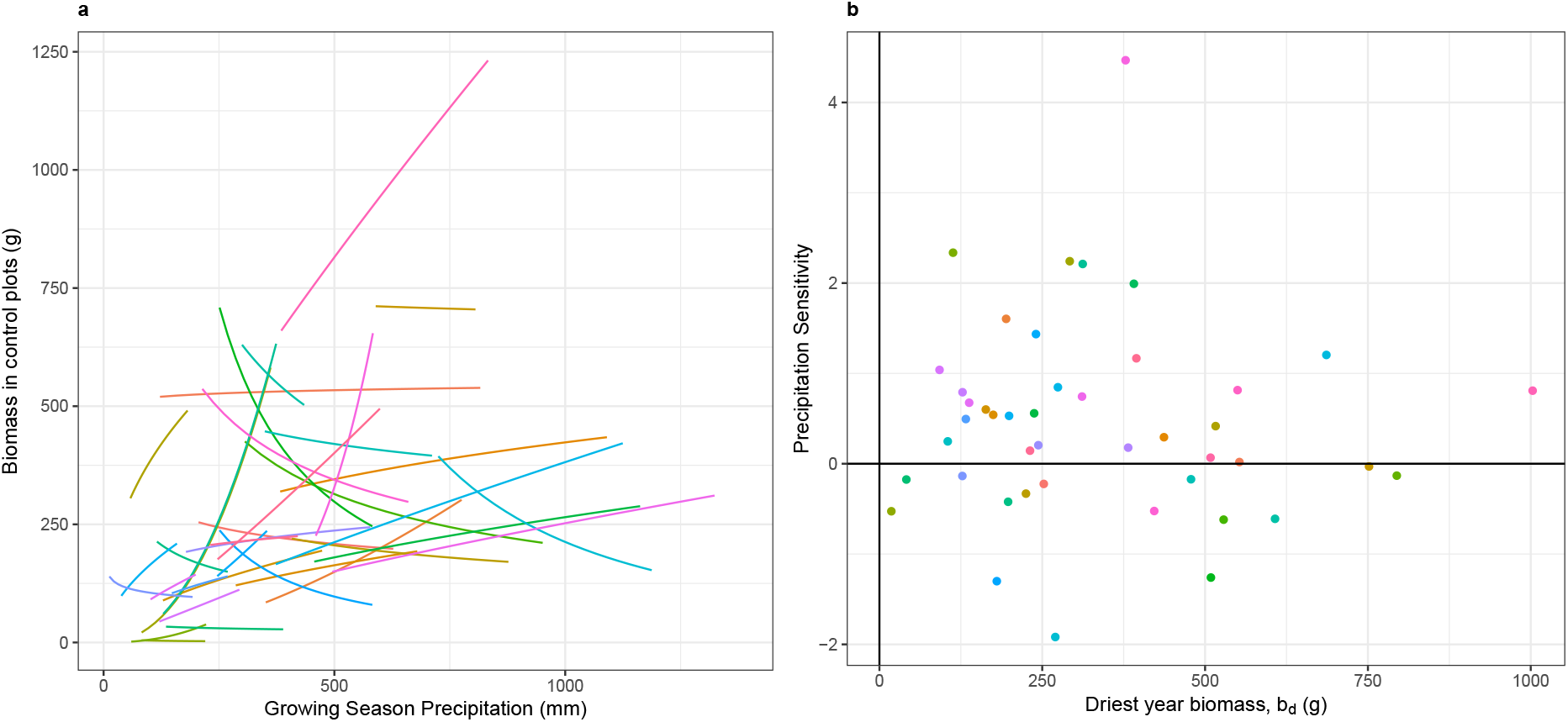
Biomass-precipitation relationships in unmanipulated plots at 44 grassland sites. Relationships were estimated by fitting linear models to *log*_2_ transformed data of both biomass and growing season precipitation (GSP). **a.** Fitted relationships in control plots in our study. **b.** Values of precipitation sensitivity S (proportional response of biomass to a change in rainfall) and biomass measured in the driest year *b_d_*.

Nutrient addition significantly increased driest year biomass (median = + 60%, Wilcoxon signed rank exact test, V = 788, p < 0.001, Figure 3b). It resulted in a marginal, but nonsignificant increase of precipitation sensitivity (median +0.08, V = 640, p = 0.09, Figure 3a). Since our study is on natural temporal variation in precipitation at each site, we also checked whether the observed effect of nutrient addition on driest year biomass depended on how extreme the driest year was at each site. We found that the nutrient effect on *b_d_* was not significantly associated with the *SPEI_g_s* value of that driest year (Appendix S1: Figure S4).

**Figure 3:**
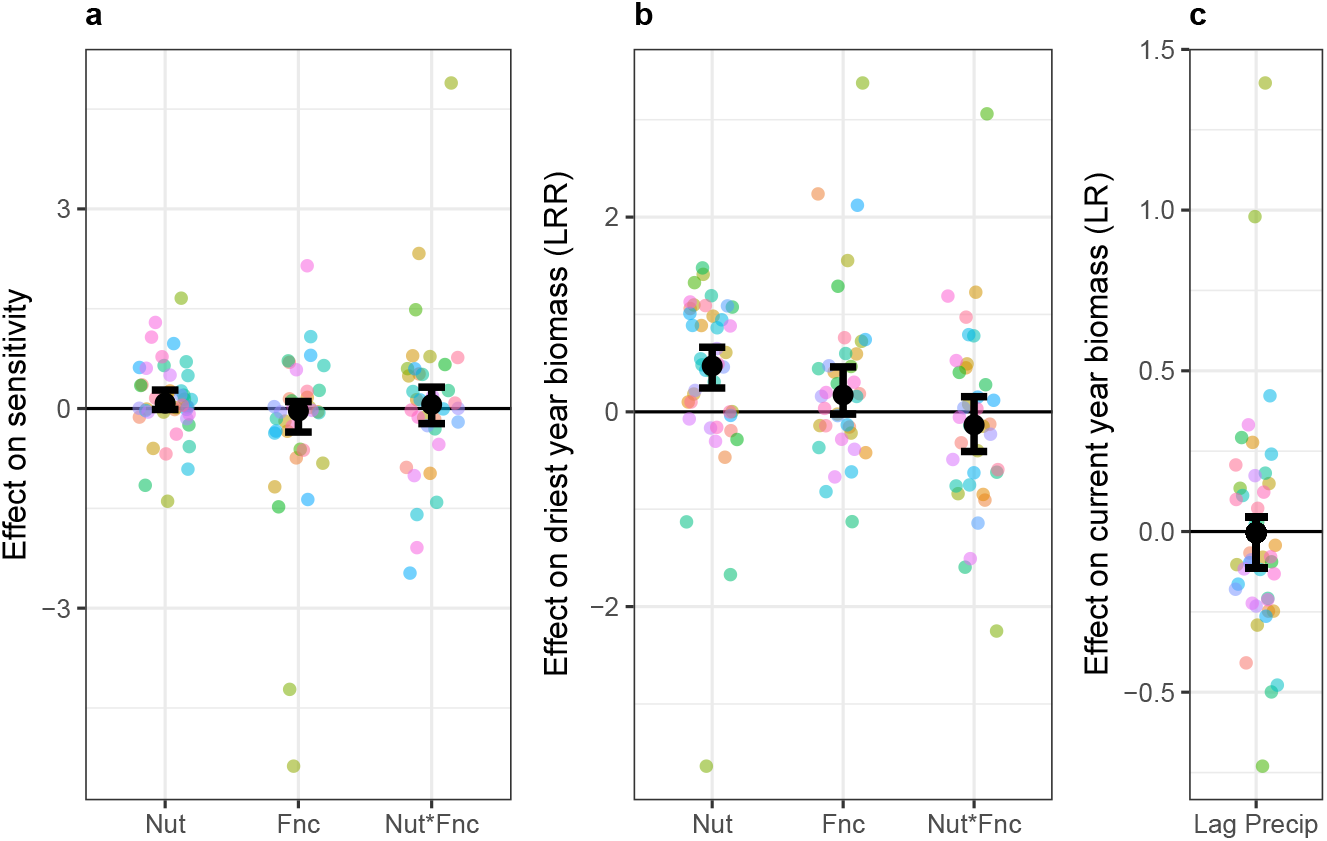
The effects of nutrient addition (Nut, n = 44), grazer exclusion (Fnc, n = 36), and their interaction (n=36) on biomass - precipitation relationships. Nut and Fnc effects are change relative to control plots. ‘Nut^*^Fnc’ denotes the interaction between the two treatments. Error bars denote 95% confidence intervals from Wilcoxon signed rank tests. **a.** The log_2_ response ratio of treatment effects on driest year biomass at each site (b_d_), **b.** The additive effects of treatments on precipitation sensitivity (*S*). **c.** The log_2_ effect of precipitation in the previous year (measured as SPEI) on current year biomass.

On average, excluding herbivores did not change the effect of fertilization on biomass-precipitation relationships. There was no interaction effect between the nutrient addition and fencing treatments on either sensitivity (median 0.06, V = 360, p = 0.68, Figure 3a) or driest year biomass (median −12%, V = 273, p = 0.35, Figure 3b).

Precipitation in the previous growing season was an important factor for explaining current year biomass at many sites. While the mean effect was not different from zero, there were sites with both strongly positive and strongly negative legacy effects (Figure 3c, Appendix S1: Figure S5). The proportion of total biomass variance explained by legacy effects ranged from 0% to 74% (median 14%, IQR = 5% to 33%). In 21 out of 44 sites (48%), previous year's water availability explained more variance in biomass than current year growing season precipitation.

### Changes with site aridity and fencing

Next, we examined how the parameter values of sensitivity and driest year biomass, and the effects of fertilization on both, varied across the aridity gradient.

Sensitivity did not significantly change from arid to mesic sites (p = 0.4, Figure 4b). Driest year biomass strongly increased from arid to mesic sites (p = 0.02), with arid sites having low biomass and mesic sites having large variation in biomass (Figure 4b).

**Figure 4:**
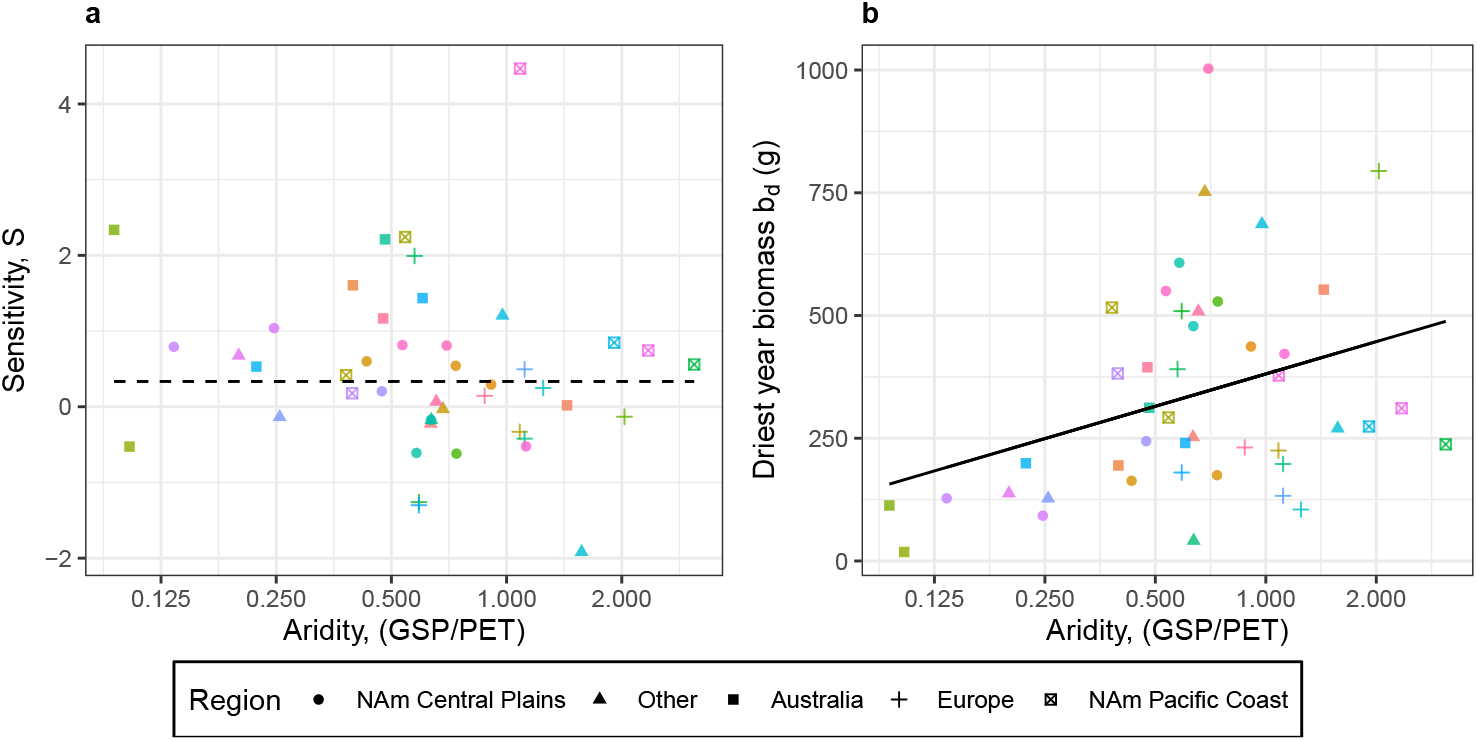
: Change of **a.** precipitation sensitivity (*S*) and **b.** driest year biomass (*b_d_*) across the
gradient of aridity among sites in this study. Aridity is measured by the ratio of mean precipitation to potential evapotranspiration over the growing season at each site. (GSP/PET) < 1 indicates dry sites where evapotranspiration exceeds precipitation, and (GSP/PET) > 1 in-dicates mesic sites with greater water availability Points denote individual sites, with shape varying by region.

Nutrient addition increased both driest year biomass and precipitation sensitivity regardless of site aridity (Figure 5b,d). The increase in driest year biomass caused by nutrient addition was constant across the aridity gradient, and was unaffected by the exclusion of grazers (Figure 5c,d). Nutrient addition also increased driest year biomass irrespective of how relatively dry that year was in comparison to the site’s climate history i.e. there was no correlation between the nutrient effect on driest year biomass and the SPEI value of that year (Appendix S1: Figure S4). Due to our effects being measured on a log_2_ scale, and biomass increasing from arid to mesic sites (Figure 4b), the constant relative nutrient effect corresponds to an absolute increase in the biomass added by nutrients in mesic versus arid sites.

**Figure 5:**
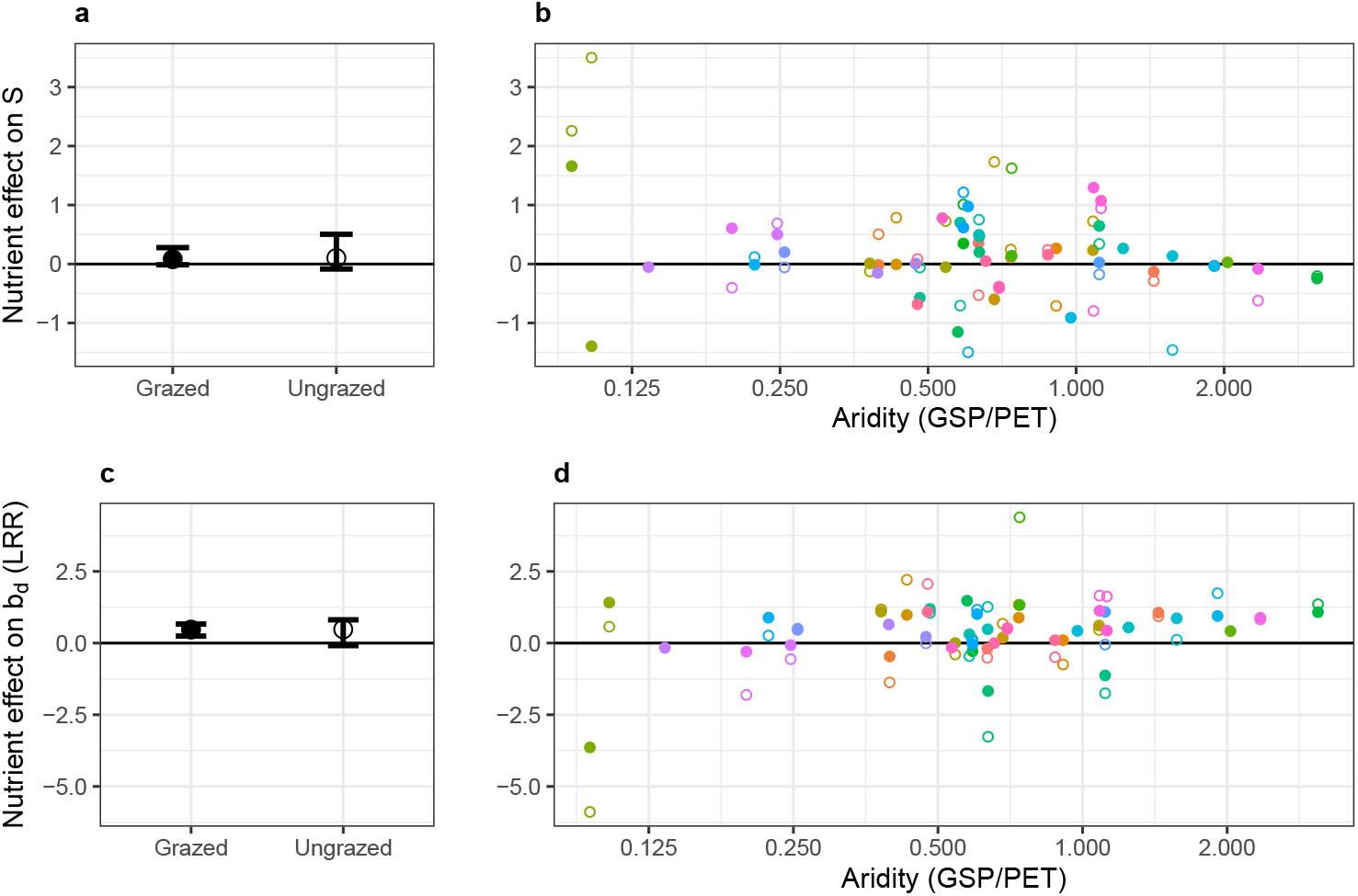
Effect of nutrient addition on sensitivity (a,b) and driest year biomass (c,d) across the aridity gradient among sites and experimental grazer exclusion in this study. Filled points denote mean effects for grazed plots at a site, whereas open points denote fenced plots. Error bars (panels a, c) show 95% confidence intervals estimated from Wilcoxon signed-rank tests. There is no significant relationship between site aridity and any of the response variables.

Nutrient effects on biomass sensitivity to precipitation had a larger variance in ungrazed plots as compared to grazed plots (F_43,35_ = 0.35, p = 0.001). Nutrient addition marginally increased sensitivity in grazed plots (mean = +0.12, Wilcoxon signed rank test V = 641, p = 0.09), and did not significantly increase sensitivity in ungrazed plots (V = 422, p = 0.16, Figure 5a). The effects of grazers on nutrient addition did not significantly change across the aridity gradient (Figure 5b). When we estimated sensitivity as the slope of linear biomass-precipitation relationships (units: *gm*^−2^*mm*^−1^), then we found that nutrient addition significantly increased sensitivity in grazed plots (V = 733, p = 0.005, Appendix S3:Figure S2). The linear slope corresponds to the increase in biomass per unit rainfall, whereas our relative sensitivity metric measures the proportional change in biomass for a change in rainfall.

## 4 Discussion

We examined the interaction between spatial and temporal variation in water availability on grassland biomass at 44 sites around the world, and how that was influenced by nutrient addition and herbivore removal. In spite of the significant heterogeneity in all responses, nutrient addition increased both driest year biomass and precipitation sensitivity across the whole range of aridity. This was contrary to our expectation that nutrients would not affect driest year biomass or sensitivity at arid sites. These findings are consistent with models of plant biomass being co-limited by nutrients and water. In almost half of our sites, the previous year's rainfall explained as much variation in biomass as current year precipitation, highlighting the importance of accounting for legacies in estimations of biomass-precipitation relationships (Sala, Gherardi, et al., 2012; Silvertown et al., 1994).

Resource co-limitation occurs when primary production is simultaneously limited by multiple resources, and shows non-additive responses to factorial resource additions (Harpole et al., 2011; Sperfeld et al., 2016). Species interactions (Danger et al., 2008), optimal foraging by plants (Rastetter and Shaver, 1992), variation in the availability of multiple resources over time (Yahdjian et al., 2011), and the dependence of plant mineral uptake on soil water availability (Everard et al., 2010; Plett et al., 2020) frequently result in biomass being co-limited by multiple resources. The degree of limitation can vary between different resources, thus leading to biomass in some ecosystems being functionally limited by just a single resource (Fay et al., 2015; Harpole et al., 2011). This underlies the expectation that arid ecosystems are fundamentally limited by water, and should be unresponsive to added nutrients in the absence of added water (Eskelinen and Harrison, 2015; Yahdjian et al., 2011). We found that nutrient addition increased driest year biomass at most sites in our study, suggesting that these grasslands are independently responsive to both water and nutrients (as per Harpole et al., 2011) Thus, our study demonstrates the ubiquity of co-limitation between water and nutrients in grasslands across a globally-representative range of aridity.

Multiple resource limitation has been proposed as a framework to explain variation in biomass sensitivity to precipitation across aridity gradients (Huxman et al., 2004). Comparison of site responses across the aridity gradient in our study directly tested and supported this framework, building on earlier meta-analyses (Hooper and Johnson, 1999; Yahdjian et al., 2011). We found that sensitivity did not significantly change between arid and mesic sites, matching other cross site observational studies (Bai et al., 2008; Hsu et al., 2012). Experimental nutrient addition increased sensitivity all across the gradient of aridity spanned by our sites, refuting our expectations from Huxman et al. (2004) that nutrient addition would have stronger effects at mesic, rather than arid sites. Many studies have sought to undertand the the effects of global change on the stability of grassland ecosystems, by examining the variation of biomass over time in different treatments (Gilbert et al., 2020; Hautier et al., 2014). In this study we focused on one aspect of that variation -how biomass is driven by inter-annual variation in rainfall. Our findings of co-limitation extend our understanding of the controls on grassland ecosystem processes across space and time, thus providing mechanistic bases for future predictions of grassland responses to global change.

Herbivores and climate can influence ecosystems by changing the availability of nutrients and water for plants (Frank et al., 2018). Our fencing treatments enabled us to evaluate the role of grazers in shaping multiple resource limitation of biomass. On average, the effect of nutrients on biomass-precipitation relationships was similar in the presence and absence of grazers. However, excluding grazers both increased and decreased the effect of nutrients on sensitivity (overall increasing the variance of nutrient effects). Excluding grazers can increase nutrient effects on sensitivity if grazers are consuming the additional biomass generated by fertilization (Gruner et al., 2008). Leaf litter can suppress plant growth, and grazers can reduce and remove standing litter, enhancing plant growth (Borer, Seabloom, et al., 2014). In such situations, excluding grazers can dampen the effects of nutrients on biomass sensitivity, which is visible in our study. Fencing also directly affected biomass-precipitation relationships. At some sites, excluding grazers increased the precipitation sensitivity of biomass, implying that grazing dampens the variation in biomass caused due to interannual variation in precipitation. At other sites, excluding grazers decreased precipitation sensitivity, implying that grazing can potentially exacerbate the effects of rainfall change (Staver et al., 2019). Thus, grazer exclusion had context-dependent effects on the interaction between nutrients and biomass precipitation relationships.

Our study is the first large scale evaluation of the effects of nutrient addition on the relationship between plant biomass and annual precipitation in grasslands, with sites spanning the range of precipitation variation experienced by global grasslands (Gilbert et al., 2020). Though there was significant variation in the shape of biomass-precipitation relationships across sites, the effects of nutrient addition supports a model of grassland biomass being co-limited by both nutrients and water, irrespective of whether a site is arid or mesic. Thus nutrient eutrophication has the potential to increase temporal biomass variation, reducing the stability of even arid grasslands (Hautier et al., 2014; Sloat et al., 2018). The presence or absence of grazers did not consistently change the resource co-limitation, although grazer effects did vary strongly among sites. Ongoing efforts of coordinating distributed experiments across regions of the world (Borer, Grace, et al., 2017; Knapp et al., 2017) will deepen our understanding of processes, patterns and contingencies driving ecosystem functions.

## 5 Acknowledgments

We would like to thank Forest Isbell and Julie Grossman for their feedback on this paper. This work was generated using data from the Nutrient Network (http://www.nutnet.org) experiment, funded at the site-scale by individual researchers. Coordination and data management have been supported by funding to E. Borer and E. Seabloom from the National Science Foundation Research Coordination Network (NSF-DEB-1042132) and Long Term Ecological Research (NSF-DEB-1234162 & DEB-1831944) programs, and the Institute on the Environment (DG-0001-13). We also thank the Minnesota Supercomputer Institute for hosting project data and the Institute on the Environment for hosting Network meetings. Author and data contributions are available in Appendix S4. USDA-ARS is an equal opportunity employer. S Bharath would like to acknowledge that the University of Minnesota Twin Cities stands on Miní Sóta Makhóčhe. Support for his research comes from this land grant institution illegally occupying the homelands of the Dakhóta Oyáte.

## Notes

### Competing Interest Statement

The authors have declared no competing interest.

