## Supplementary materials for "Multiple resource limitations explain biomass-precipitation relationships in grasslands"

**Siddharth Bharath et al. Multiple resource limitation explains biomass-precipitation relationships in global grasslands. Supplementary Materials**

#### Appendix S1

##### Details of model fits

###### Figure S1: Model outputs graphed as ANPP~Precip relations

This graph shows the raw data and the fitted models for each treatment at each site. Model fits are accounting for the lagged effects of previous years precipitation on current year biomass.

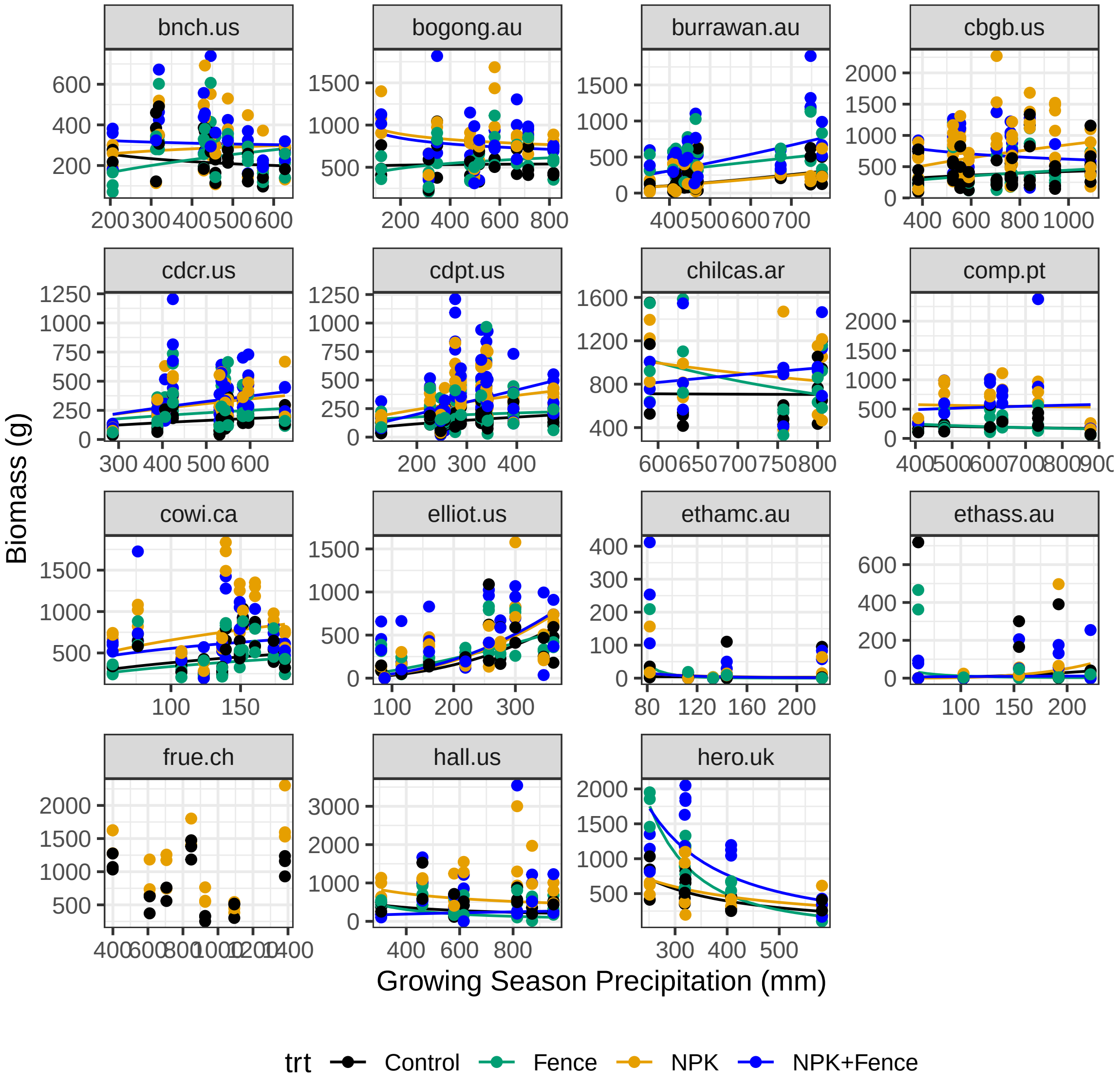

Figure S1 continued

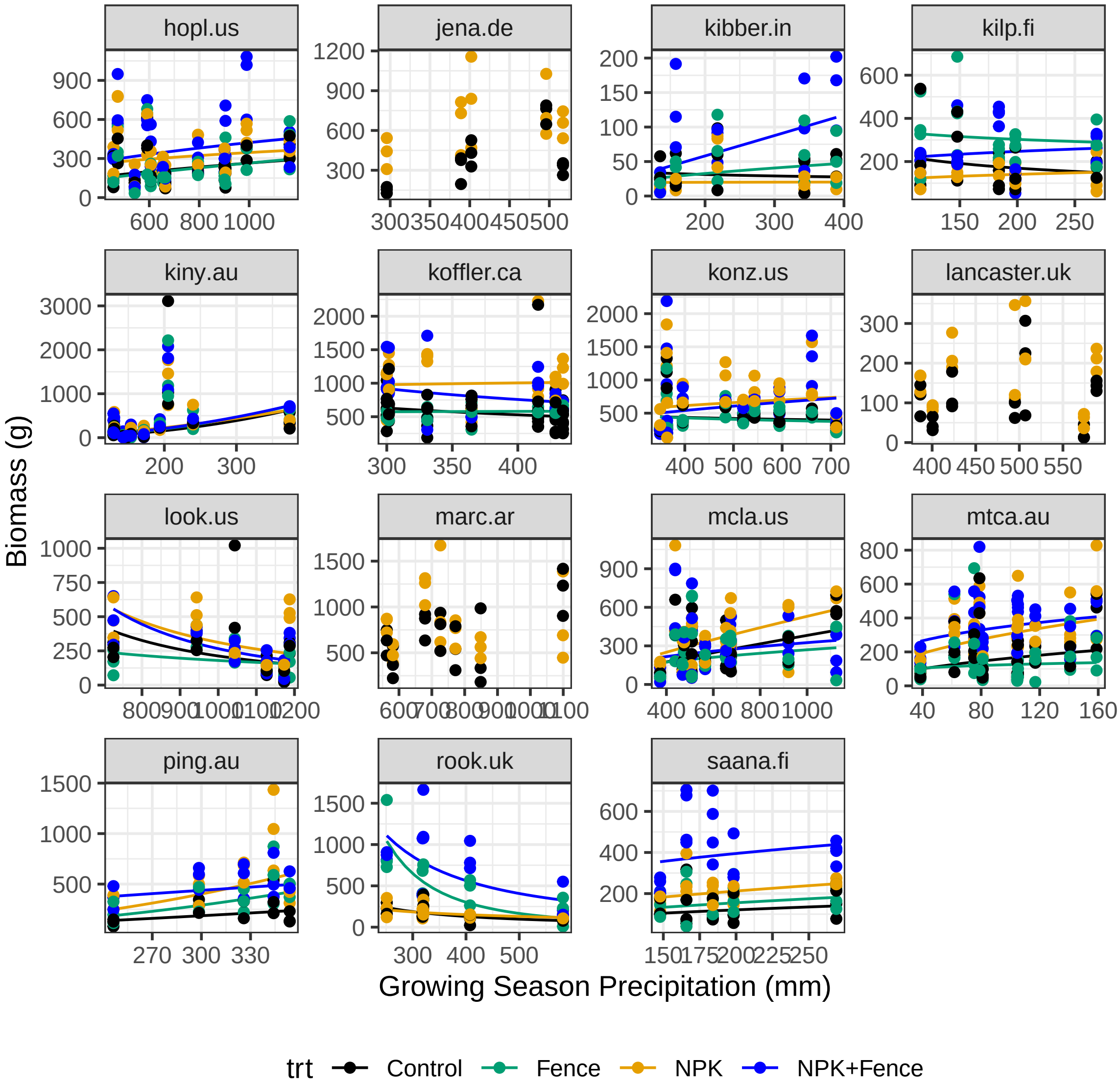

Figure S1 continued

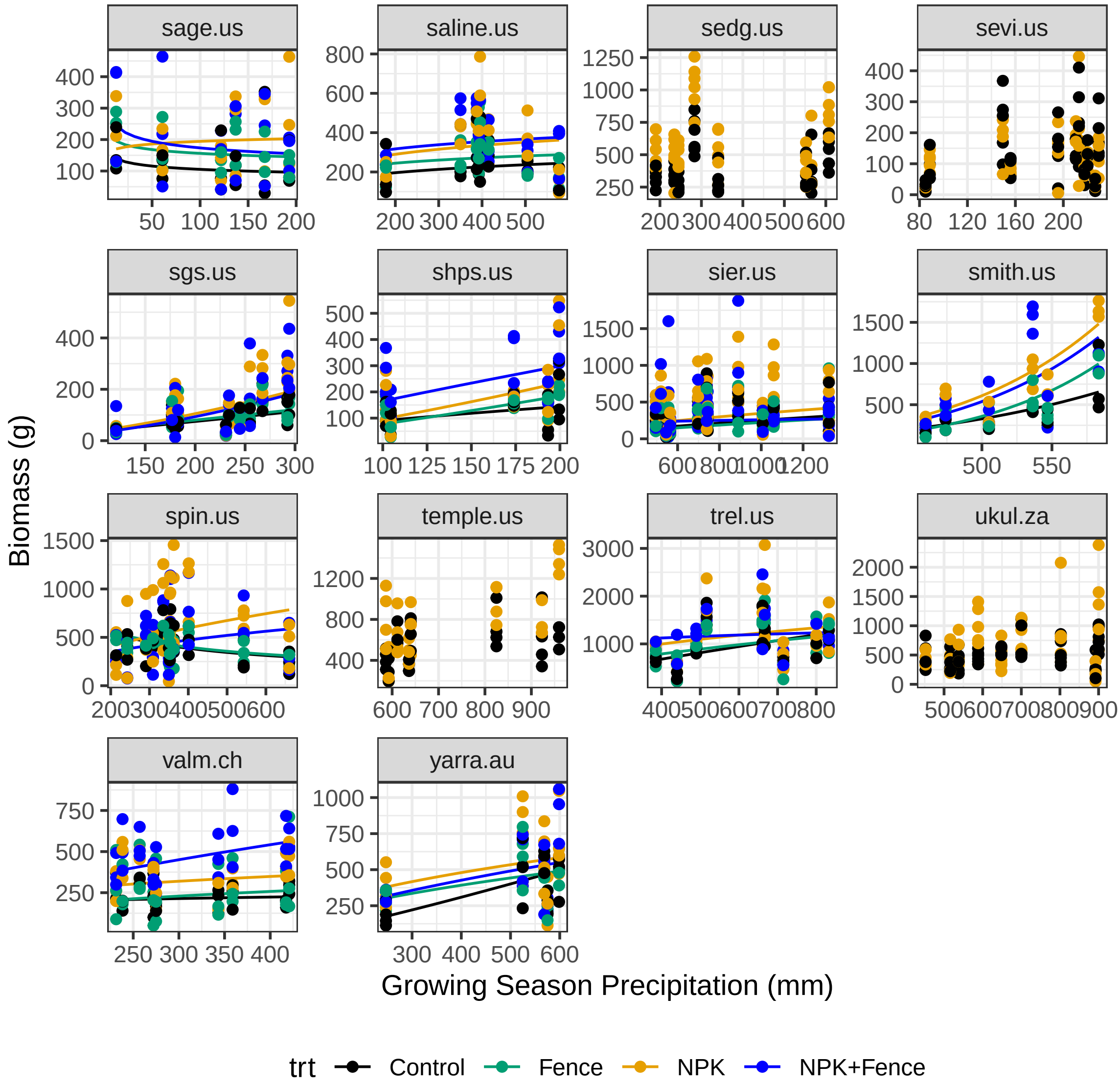

**Table S2: Sensitivity values and treatment effects for each site**

| Site Code | Latitude | Longitude | Sensitivity (g/mm) | Nut | Fence | Nut*Fnc |
|---|---|---|---|---|---|---|
| bnch.us | 44.28 | -121.97 | -0.22 | 0.36 | 0.19 | -0.88 |
| bogong.au | -36.87 | 147.25 | 0.02 | -0.13 | -0.16 | -0.16 |
| burrawan.au | -27.73 | 151.14 | 1.60 | -0.02 | 2.24 | 0.52 |
| cbgb.us | 41.79 | -93.39 | 0.29 | 0.26 | -0.42 | -0.97 |
| cdcr.us | 45.42 | -93.21 | 0.54 | 0.11 | 0.41 | 0.13 |
| cdpt.us | 41.20 | -101.63 | 0.60 | -0.01 | 0.59 | 0.79 |
| chilcas.ar | -36.28 | -58.27 | -0.03 | -0.60 | -0.02 | 2.33 |
| comp.pt | 38.83 | -8.79 | -0.33 | 0.23 | -0.14 | 0.49 |
| cowi.ca | 48.81 | -123.63 | 0.42 | 0.01 | -0.22 | -0.14 |
| elliot.us | 32.88 | -117.05 | 2.24 | -0.05 | 1.55 | 0.78 |
| ethamc.au | -23.76 | 138.47 | -0.53 | -1.39 | 0.72 | 4.89 |
| ethass.au | -23.64 | 138.40 | 2.34 | 1.66 | 3.38 | 0.60 |
| frue.ch | 47.11 | 8.54 | -0.13 | 0.03 | NA | NA |
| hall.us | 36.87 | -86.70 | -0.62 | 0.14 | 0.47 | 1.48 |
| hero.uk | 51.41 | -0.64 | -1.26 | 0.35 | 1.29 | 0.66 |
| hopl.us | 39.01 | -123.06 | 0.56 | -0.25 | 0.29 | 0.04 |
| jena.de | 50.93 | 11.53 | 1.99 | -1.15 | NA | NA |
| kibber.in | 32.32 | 78.01 | -0.17 | 0.20 | -1.13 | 0.27 |
| kilp.fi | 69.06 | 20.87 | -0.42 | 0.65 | 0.44 | -0.31 |
| kiny.au | -36.20 | 143.75 | 2.21 | -0.57 | 0.60 | 0.51 |
| koffler.ca | 44.02 | -79.54 | -0.61 | 0.70 | -0.37 | -1.41 |
| konz.us | 39.07 | -96.58 | -0.17 | 0.50 | 0.16 | 0.26 |
| lancaster.uk | 53.99 | -2.63 | 0.25 | 0.26 | NA | NA |
| look.us | 44.21 | -122.13 | -1.92 | 0.14 | -0.82 | -1.59 |
| marc.ar | -37.72 | -57.42 | 1.20 | -0.91 | NA | NA |
| mcla.us | 38.86 | -122.41 | 0.85 | -0.03 | -0.62 | -0.01 |
| mtca.au | -31.78 | 117.61 | 0.53 | -0.01 | -0.13 | 0.13 |
| ping.au | -32.50 | 116.97 | 1.44 | 0.98 | 0.74 | -2.47 |
| rook.uk | 51.41 | -0.64 | -1.30 | 0.62 | 2.12 | 0.60 |
| saana.fi | 69.04 | 20.84 | 0.49 | 0.03 | -0.04 | -0.20 |
| sage.us | 39.43 | -120.24 | -0.14 | 0.20 | 0.47 | -0.26 |
| saline.us | 39.05 | -99.10 | 0.21 | 0.00 | 0.16 | 0.00 |

| Site Code | Latitude | Longitude | Sensitivity (g/mm) | Nut | Fence | Nut*Fnc |
|---|---|---|---|---|---|---|
| sedg.us | 34.70 | -120.02 | 0.18 | -0.15 | NA | NA |
| sevi.us | 34.36 | -106.69 | 0.79 | -0.05 | NA | NA |
| sgs.us | 40.82 | -104.77 | 1.04 | 0.50 | -0.38 | 0.19 |
| shps.us | 44.24 | -112.20 | 0.68 | 0.61 | -0.67 | -1.01 |
| sier.us | 39.24 | -121.28 | 0.74 | -0.08 | -0.28 | -0.54 |
| smith.us | 48.21 | -122.62 | 4.47 | 1.29 | 0.20 | -2.09 |
| spin.us | 38.14 | -84.50 | -0.52 | 1.07 | 0.30 | -0.13 |
| temple.us | 31.04 | -97.35 | 0.81 | 0.78 | NA | NA |
| trel.us | 40.08 | -88.83 | 0.81 | -0.38 | 0.04 | -0.02 |
| ukul.za | -29.67 | 30.40 | 0.07 | 0.05 | NA | NA |
| valm.ch | 46.63 | 10.37 | 0.14 | 0.16 | -0.14 | 0.08 |
| yarra.au | -33.61 | 150.73 | 1.17 | -0.68 | 0.76 | 0.77 |

**Table S3: Driest year biomass values and treatment effects for each site**

| site_code | Biomassd (g) | Nut (LRR) | Fnc (LRR) | Nut*Fnc (LRR) | Lagged PPT (LR) |
|---|---:|---:|---:|---:|---:|
| bnch.us | 7.98 | -0.19 | 0.19 | -0.32 | -0.41 |
| bogong.au | 9.11 | 1.06 | -0.16 | -0.13 | -0.07 |
| burrawan.au | 7.60 | -0.47 | 2.24 | -0.91 | -0.25 |
| cbgb.us | 8.77 | 0.10 | -0.42 | -0.85 | -0.04 |
| cdcr.us | 7.45 | 0.89 | 0.41 | 0.45 | 0.28 |
| cdpt.us | 7.35 | 0.98 | 0.59 | 1.23 | -0.25 |
| chilcas.ar | 9.55 | 0.18 | -0.02 | 0.49 | -0.08 |
| comp.pt | 7.81 | 0.61 | -0.14 | -0.15 | 0.15 |
| cowi.ca | 9.01 | 1.10 | -0.22 | 0.08 | -0.29 |
| elliot.us | 8.19 | 0.00 | 1.55 | -0.40 | -0.10 |
| ethamc.au | 4.20 | 1.41 | 0.72 | -0.84 | 1.40 |
| ethass.au | 6.82 | -3.64 | 3.38 | -2.25 | 0.98 |
| frue.ch | 9.63 | 0.42 | NA | NA | 0.13 |
| hall.us | 9.05 | 1.33 | 0.47 | 3.06 | -0.73 |
| hero.uk | 8.99 | -0.28 | 1.29 | 0.41 | -0.09 |
| hopl.us | 7.89 | 1.08 | 0.29 | 0.28 | 0.29 |
| jena.de | 8.61 | 1.48 | NA | NA | 0.03 |
| kibber.in | 5.37 | -1.67 | -1.13 | -1.60 | -0.50 |
| kilp.fi | 7.63 | -1.13 | 0.44 | -0.62 | -0.21 |
| kiny.au | 8.29 | 1.19 | 0.60 | -0.14 | 0.18 |
| koffler.ca | 9.25 | 0.31 | -0.37 | -0.76 | 0.02 |
| konz.us | 8.90 | 0.48 | 0.16 | 0.78 | 0.11 |
| lancaster.uk | 6.71 | 0.54 | NA | NA | -0.48 |
| look.us | 8.08 | 0.86 | -0.82 | -0.75 | -0.12 |
| marc.ar | 9.42 | 0.43 | NA | NA | 0.24 |
| mcla.us | 8.10 | 0.95 | -0.62 | 0.79 | 0.42 |
| mtca.au | 7.64 | 0.89 | -0.13 | -0.63 | -0.16 |
| ping.au | 7.91 | 1.01 | 0.74 | 0.15 | -0.26 |
| rook.uk | 7.49 | -0.04 | 2.12 | 0.12 | -0.10 |
| saana.fi | 7.05 | 1.09 | -0.04 | -1.14 | 0.00 |
| sage.us | 6.99 | 0.46 | 0.47 | 0.04 | -0.18 |
| saline.us | 7.93 | 0.22 | 0.16 | -0.23 | -0.09 |

| site_code | Biomassd (g) | Nut (LRR) | Fnc (LRR) | Nut*Fnc (LRR) | Lagged PPT (LR) |
|---|---|---|---|---|---|
| sedg.us | 8.58 | 0.65 | NA | NA | -0.23 |
| sevi.us | 7.00 | -0.17 | NA | NA | 0.17 |
| sgs.us | 6.53 | -0.07 | -0.38 | -0.49 | -0.12 |
| shps.us | 7.11 | -0.30 | -0.67 | -1.51 | -0.22 |
| sier.us | 8.28 | 0.88 | -0.28 | -0.06 | 0.33 |
| smith.us | 8.56 | 1.13 | 0.20 | 0.53 | -0.21 |
| spin.us | 8.72 | 0.43 | 0.30 | 1.19 | -0.13 |
| temple.us | 9.10 | -0.16 | NA | NA | -0.08 |
| trel.us | 9.97 | 0.49 | 0.04 | 0.03 | 0.12 |
| ukul.za | 8.99 | 0.00 | NA | NA | 0.10 |
| valm.ch | 7.85 | 0.10 | -0.14 | -0.59 | 0.07 |
| yarra.au | 8.62 | 1.09 | 0.76 | 0.97 | 0.21 |

**Figure S4: Nutrient effects on driest year biomass compared to year aridity**

Does the effect size of nutrient addition on driest year biomass depend on how dry that year was? Points denote individual sites, with the x axis value being the growing season SPEI value of the driest year at that site.

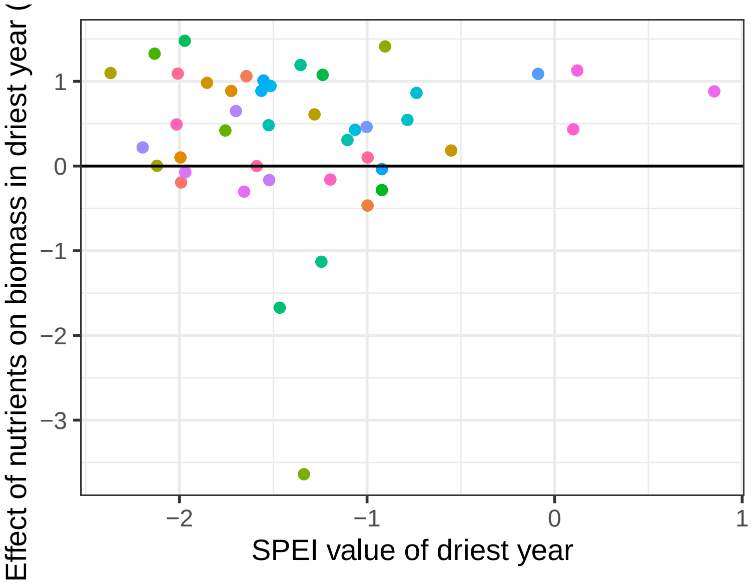

**Figure S5: Lagged precipitation effects**

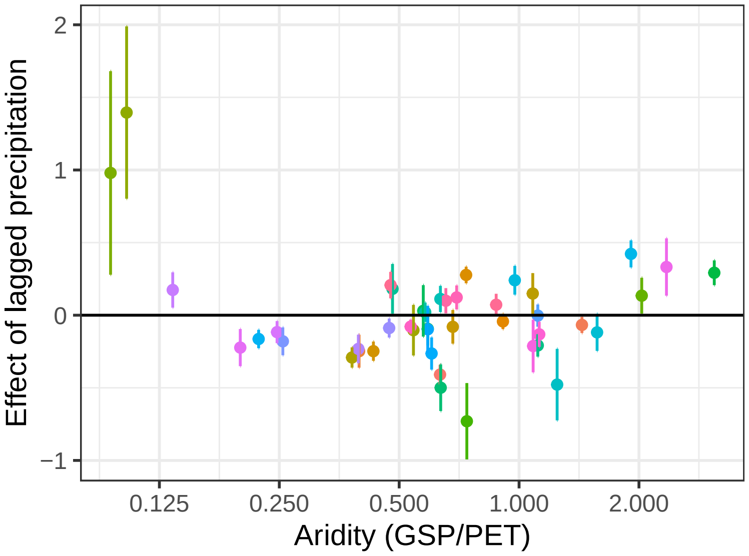


Estimation of the effect of previous year's water availability (quantified as the growing season SPEI) on current year's biomass, and it's relationship to site aridity. Error bars denote standard errors around the mean.

**Figure S6: Joint treatment effects on sensitivity and $b_d$**

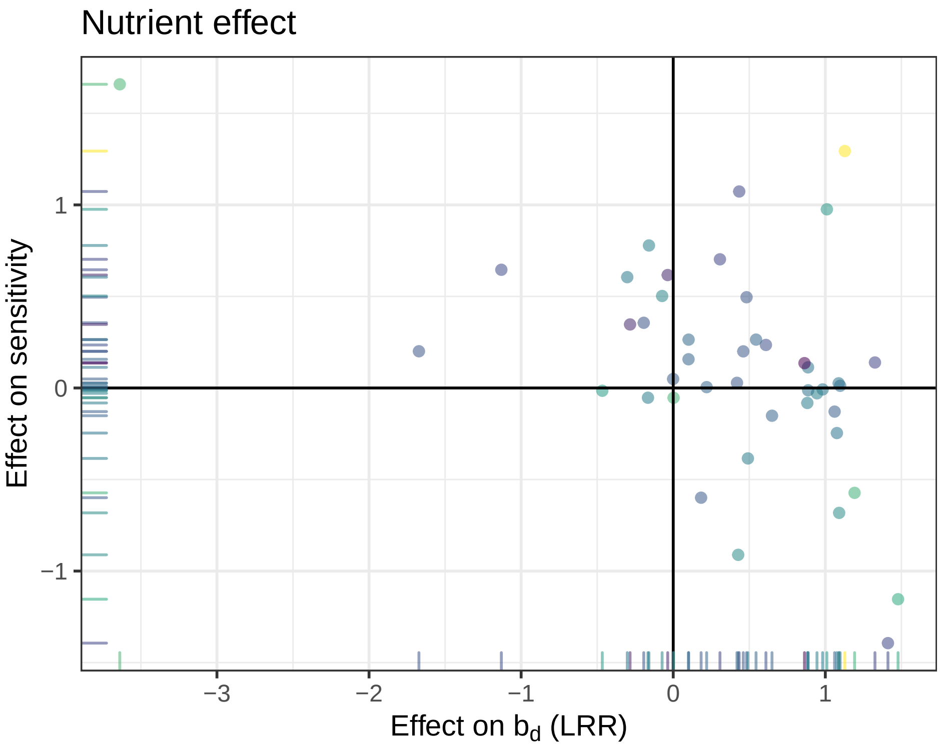


We can use this graph of nutrient effects on $b_d$ and $S$ to characterize colimitation between water and nutrients as falling into the different categories described in Box 1 of Harpole, W. S., J. T. Ngai, E. E. Cleland, E. W. Seabloom, E. T. Borer, M. E. S. Bracken, J. J. Elser, D. S. Gruner, H. Hillebrand, J. B. Shurin, and J. E. Smith. 2011. Nutrient co-limitation of primary producer communities: Community co-limitation. Ecology Letters 14:852–862.

- Simultaneous colimitation -
  - Sensitivity = 0
  - Nut effect on $b_d$ = 0
  - Nut effect on $S$ = +
- Independent colimitation -
  - Sensitivity = +
  - Nut effect on $b_d$ = +

- Nut effect on $S$ = + for superadditive | 0 for additive | - for sub-additive

- Serial limitation of first water and then nutrient -
  - Sensitivity = +

  - Nut effect on $b_d$ = 0

  - Nut effect on $S$ = +

- Serial limitation of first nutrient and then water -
  - Sensitivity = 0

  - Nut effect on $b_d$ = +

  - Nut effect on $S$ = +

**Siddharth Bharath et al. Multiple resource limitation explains biomass-precipitation relationships in global grasslands. Supplementary Materials**

#### Appendix S2

**Site information and Weather data sources**

#### Table S1 : Site experiment deviations

The standard protocol for fencing followed by most sites in this analysis is : Fences were 230 cm tall with the lower 90 cm surrounded by 1-cm woven wire mesh. An additional 30-cm outward-facing flange was stapled to the ground to exclude digging animals (for example, rabbits, voles), although not fully subterranean animals (for example, gophers, moles). Four strands of barbless wire were strung at equal vertical distances above the wire mesh.

Below is a list of sites with variations in fencing treatment different from the standard protocols. -

cdpt.us | Central plains (North America) | Five feet of 10cm mesh cattle panels, with hardware cloth up to 50cm from ground level.

hall.us | Central plains (North America) | Similar to NutNet standard with two modifcations: 1/4-inch hardware cloth instead of 1/2-inch, and 5’ fences instead of 7’ fences.

shps.us | Montane West (North America) | Similar to NutNet standard but top strand at 1.2 m

spin.us | Central plains (North America) | Similar to NutNet standard with two modifcations: 1/4-inch hardware cloth instead of 1/2-inch, and 5’ fences instead of 7’ fences.

valm.ch | Europe | 2.7 m wooden poles (25 cm diameter) driven 70 cm into ground, 3 m apart, covered with 5 cm square mesh to 2 m high and with exta cabling and supports to prevent snow damage. Fences enclose 6 m x 7 m area.

**Table S2 : Weather data sources**

| Site Code | Growing Season | Data Duration | Weather source | Distance |
|---|---|---|---|---|
| bnch.us | 4-8 | 53 \| 11 | USC00350652, USS0021E07S | 8.3 |
| bogong.au | 10-1 | 24 \| 10 | ASN00083084 | 2.0 |
| burrawan.au | 10-5 | 51 \| 11 | ASN00041025, ASN00041062 | 19.4 |
| cbgb.us | 5-10 | 42 \| 10 | USC00130203, USW00054902 | 25.3 |
| cdcr.us | 4-8 | 53 \| 11 | USC00211227, USC00212881 | 20.4 |
| cdpt.us | 4-7 | 22 \| 11 | Keystone 3W Beta | 1.7 |
| chilcas.ar | 8-3 | 8 \| 5 | Dolores | 48.8 |
| comp.pt | 10-5 | 10 \| 6 | Evora | 81.9 |
| cowi.ca | 4-7 | 44 \| 11 | CA001016995 | 10.2 |
| elliot.us | 11-4 | 52 \| 10 | USW00003131, USW00093107 | 9.1 |
| ethamc.au | 5-4 | 24 \| 5 | Main Camp North | 5.2 |
| ethass.au | 5-4 | 24 \| 5 | South Site | 0.0 |
| frue.ch | 4-9 | 13 \| 7 | Fruebuel | 0.4 |
| hall.us | 4-9 | 43 \| 7 | USC00157049 | 16.8 |
| hero.uk | 4-10 | 32 \| 5 | Silwood | 0.3 |
| hopl.us | 11-4 | 19 \| 12 | USW00023275 | 17.5 |
| jena.de | 3-10 | 46 \| 5 | GM000004204 | 3.6 |
| kibber.in | 5-8 | 38 \| 5 | NA | NA |
| kilp.fi | 6-8 | 41 \| 5 | FIE00146618 | 2.3 |
| kiny.au | 5-10 | 49 \| 10 | ASN00080002, ASN00080014 | 13.2 |
| koffler.ca | 4-8 | 14 \| 7 | CAW00064757 | 30.4 |
| konz.us | 5-9 | 22 \| 10 | USW00003936, USW00053974 | 7.6 |
| lancaster.uk | 3-8 | 51 \| 7 | Hazelrigg | 25.5 |
| look.us | 3-8 | 24 \| 6 | UPLMET | 3.2 |
| marc.ar | 4-12 | 48 \| 7 | Mar del Plata | 0.0 |
| mcla.us | 11-4 | 33 \| 12 | Knoxville creek | 1.0 |
| mtca.au | 8-10 | 52 \| 10 | ASN00010044, ASN00010121 | 19.1 |
| ping.au | 4-10 | 47 \| 5 | ASN00010524, ASN00010626 | 12.6 |
| rook.uk | 4-10 | 32 \| 5 | Silwood | 0.6 |
| saana.fi | 6-8 | 41 \| 5 | FIE00146618 | 3.3 |
| sage.us | 4-7 | 41 \| 6 | USS0020K04S | 4.8 |
| saline.us | 5-9 | 52 \| 8 | US1KSEL0001, USC00145628 | 19.5 |

| Site Code | Growing Season | Data Duration | Weather source | Distance |
|---|---|---|---|---|
| sedg.us | 11-7 | 49 \| 8 | USC00041253 | 13.5 |
| sevi.us | 4-11 | 28 \| 11 | LTER_Deepwell | 0.0 |
| sgs.us | 4-8 | 16 \| 11 | USW00094074 | 1.5 |
| shps.us | 4-9 | 53 \| 5 | USC00102707 | 0.2 |
| sier.us | 11-4 | 51 \| 12 | USC00043800 | 1.5 |
| smith.us | 10-6 | 49 \| 6 | USC00451783 | 5.0 |
| spin.us | 3-5 | 50 \| 12 | USC00153194 | 9.8 |
| temple.us | 3-10 | 51 \| 8 | USC00418646 | 17.1 |
| trel.us | 4-9 | 52 \| 9 | USC00111655, USC00111743 | 12.6 |
| ukul.za | 9-4 | 41 \| 9 | Ukulinga | 0.0 |
| valm.ch | 6-8 | 16 \| 9 | Buffalora | 8.3 |
| yarra.au | 9-3 | 25 \| 5 | ASN00067105 | 4.4 |

'Site Code' has the code name for each site followed by the number of replicate blocks at that site in parentheses.

'Growing season' lists the start and end month of the growing season considered for the site, separated by a hyphen. 10-5 implies a growing season that starts in October and ends in May.

'Data duration' lists the number of years of weather data, and biomass data available for each site, separated by a comma.

'Distance' denotes the distance of the weather source from the site. If there is more than one weather source, the average distance is reported. Weather sources that start with US, AS, GM, FI, CA are station codes from the Global Historical Climatology Network daily database (https://www.ncdc.noaa.gov/data-access/land-based-station-data/land-based-datasets/global-historical-climatology-network-ghcn). Weather source for site kibber.in is monthly gridded data from CRU TS v4.03, running from 1981 to 2018. All other weather data are from weather stations at or near the site, provided by individual site PIs.

**Siddharth Bharath et al. Multiple resource limitation explains biomass-precipitation relationships in global grasslands. Supplementary Materials**

### Appendix S3

##### Results from linear production-precipitation relationships

All the results concerning biomass sensitivity to precipitation reported in the paper are calculated by fitting linear models to $log_2$ transformed data of biomass and growing season precipitation. Here we report the same set of comparisons made on linear models fitted directly to the raw data of biomass and precipitation. This has the disadvantage of not allowing nonlinear fits, and also results in predicting negative biomass values for some sites.

We fit linear mixed effects models of the following form at each site:

$$Biomass_t \sim GSP_t \times Nutrients \times Fencing + SPEI_{t-1}$$

Peak biomass in each year was the response variable. Predictors included GSP, fencing, nutrient addition, and all interactions between the three of these, allowing both slope and intercept to vary for each treatment at a site. The SPEI is a normalized metric of water availability in a given year relative to the precipitation and temperature history of the site. This metric is positive if the previous year was wetter than the mean, and negative if it was drier than the mean. We also included a random effect for blocks within sites, to correctly account for the design of our experiment.

By this method, estimated values of sensitivity range from -1.4833639, 3.9987891.

#### Figure S1 : Linear relationships fitted at site scale

Relationship of biomass to precipitation in unmanipulated plots at 44 grassland sites. Relationships were estimated by fitting linear models to data of both biomass and growing season precipitation (GSP).

a. Fitted relationships in control plots in our study.

b. Values of precipitation sensitivity $S$ (response of biomass to a change in rainfall) and biomass measured in the driest year $b_d$.

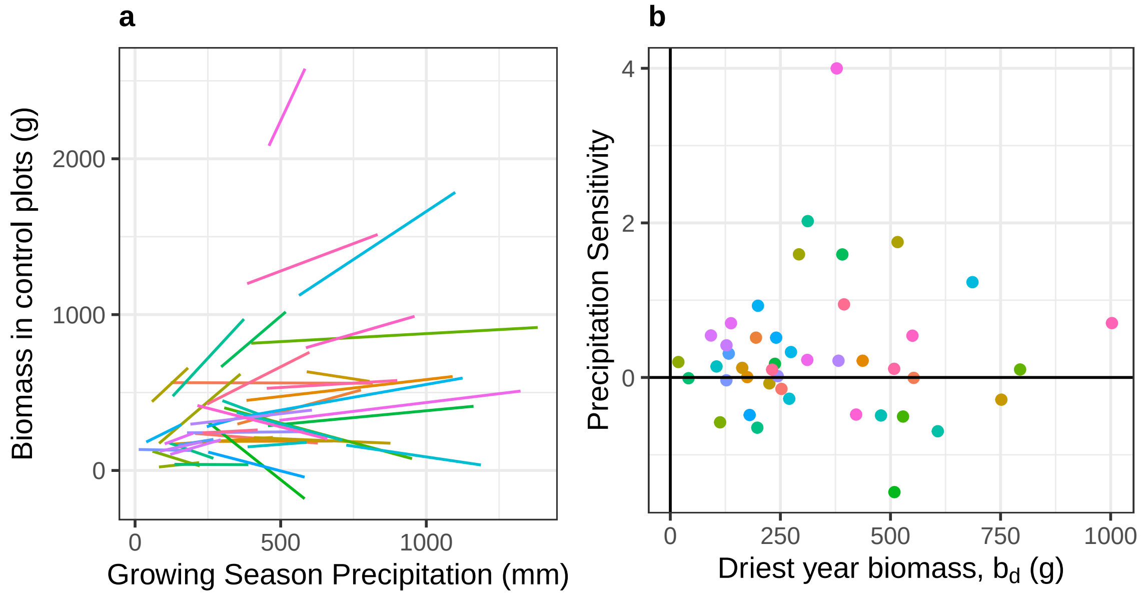


This is a complement to Figure 1 of the manuscript, just with precipitation sensitivity estimated through linear models, instead of log-log fitting.

**Figure S3 Change in sensitivity across the aridity gradient**

Change of precipitation sensitivity ($S$, units of $gm^{-2}mm^{-1}$) across the gradient of aridity among sites in this study. Aridity is measured by the ratio of mean precipitation to potential evapotranspiration over the growing season at that site, which increases from arid to mesic sites. Points denote individual sites, with shape varying by region. Sites from 4 regions with fewer than 5 sites each are labeled as 'Other'. The dotted line shows the weighted mean of sensitivity values, weighted by the inverse of the standard error of the slope measured at each site.

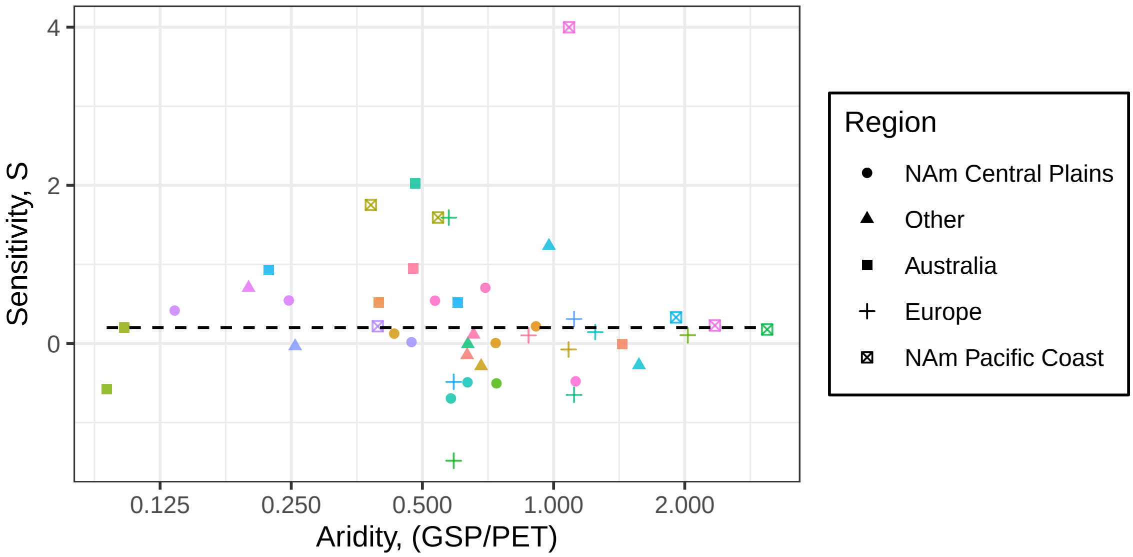

#### Figure S4 Effect of Nutrient addition on linear slope estimates

Effect of nutrient addition on sensitivity across the aridity gradient among sites and experimental grazer exclusion in this study. Filled points denote mean effects for grazed plots at a site, whereas open points denote fenced plots. Error bars (panel a) show 95% confidence intervals calculated from Wilcoxon signed-rank tests. There is no significant relationship between site aridity and any of the response variables.

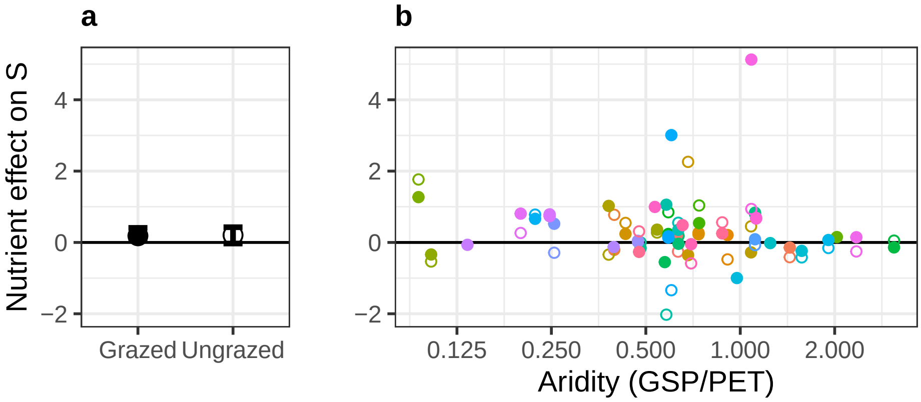


Nutrient addition significantly increases precipitation sensitivity (Wilcoxon signed rank exact test, p = 0.0047971) in the ungrazed condition, but not in the grazed condition (p = 0.0797356)

**Siddharth Bharath et al. Multiple resource limitation explains biomass-precipitation relationships in global grasslands. Supplementary Materials**

### Appendix S4 – Author and data contributions

#### Author contribution checklist –

as per the Nutrient Network co-authorship policies - https://nutnet.org/index.php/authorship

| Name | Developed and framed research question(s) | Analyzed data | Contributed to data analyses | Wrote the paper | Contributed to paper writing | Site coordinator (data collection) | Nutrient Network coordinator |
|---|---|---|---|---|---|---|---|
| Siddharth Bharath | x | x | | x | | | |
| Peter B. Adler | | | x | | x | x | |
| Philip A Fay | x | | x | | x | x | |
| Eric W. Seabloom | x | | | | x | x | x |
| Yann Hautier | | | x | | x | x | |
| Lori Biederman | | | | | x | x | |
| Miguel N. Bugalho | | | | | x | x | |
| Maria Caldeira | | | | | x | x | |
| Anu Eskelinen | | | | | x | x | |
| Johannes M H Knops | | | | | x | x | |
| John Morgan | | | | | x | x | |
| Sally A Power | | | | | x | x | |
| McCulley Rebecca | | | | | x | x | |
| Anita C. Risch | | | | | x | x | |
| Martin Schuetz | | | | | x | x | |
| Carly J. Stevens | | | | | x | x | |
| Ohlert Timothy | | | | | x | x | |
| Risto Virtanen | | | | | x | x | |
| Elizabeth Borer | x | | | | x | x | x |

**Data contributors -**
The following scientists contributed data to this study as principal investigators of their NutNet site. This cooperative research would not have been possible without their sustained efforts and contributions.

1. Bogong (bogong.au) - John Morgan, Joslin Moore
2. Bunchgrass (Andrews LTER) (bnch.us) - Eric Seabloom, Elizabeth Borer
3. Burrawan (burrawan.au) - Jennifer Firn
4. Cedar Creek LTER (cdcr.us) - Eric Seabloom, Elizabeth Borer, W. Stanley Harpole
5. Cedar Point Biological Station (cdpt.us) - Johannes Knops, George Wheeler
6. Chichaqua Bottoms (cbgb.us) - W. Stanley Harpole, Lori Biederman, Kirsten Hofmockel, Lauren Sullivan
7. Companhia das Lezirias (comp.pt) - Maria Caldeira, Miguel Bugalho
8. Cowichan (cowi.ca) - Andrew MacDougall
9. Elliott Chaparral (elliot.us) - Elsa Cleland
10. Ethabuka (Main Camp) (ethamc.au) - Glenda Wardle
11. Ethabuka (South Site) (ethass.au) - Glenda Wardle
12. Fruebuel (frue.ch) - Andy Hector, Yann Hautier, Sabine Güsewell
13. Hall's Prairie (hall.us) - Rebecca McCulley, Jim Nelson
14. Heronsbrook (Silwood Park) (hero.uk) - Mick Crawley
15. Hopland REC (hopl.us) - Eric Seabloom, Elizabeth Borer, W. Harpole
16. JeNut (jena.de) - Anne Ebeling, Christiane Roscher
17. Kibber (Spiti) (kibber.in) - Mahesh Sankaran
18. Kilpisjärvi (kilp.fi) - Anu Eskelinen, Risto Virtanen
19. Kinypanial (kiny.au) - John Morgan
20. Koffler Scientific Reserve at Joker's Hill (koffler.ca) - Arthur Weiss, Marc Cadotte
21. Konza LTER (konz.us) - Melinda Smith, Kimberly Komatsu
22. Lancaster (lancaster.uk) - Carly Stevens
23. Las Chilcas (chilcas.ar) - Laura Yahdjian, Enrique Chaneton
24. Lookout (Andrews LTER) (look.us) - Eric Seabloom, Elizabeth Borer
25. Mar Chiquita (marc.ar) - Pedro Daleo, Juan Alberti
26. Mclaughlin UCNRS (mcla.us) - Eric Seabloom, Elizabeth Borer, W. Harpole
27. Mt. Caroline (mtca.au) - Suzanne Prober
28. Pingelly Paddock (ping.au) - Jodi Price, Rachel Standish
29. Rookery (Silwood Park) (rook.uk) - Mick Crawley
30. Saana (saana.fi) - Anu Eskelinen, Risto Virtanen
31. Sagehen Creek UCNRS (sage.us) - Daniel Gruner, Louie Yang
32. Saline Experimental Range (saline.us) - Melinda Smith, Kimberly Komatsu
33. Sedgwick Reserve UCNRS (sedg.us) - Carla D'Antonio, W. Harpole, Elizabeth Borer, Eric Seabloom
34. Sevilleta LTER (sevi.us) - Scott Collins, Timothy Ohlert
35. Sheep Experimental Station (shps.us) - Peter Adler
36. Shortgrass Steppe LTER (sgs.us) - Cynthia Brown, Julia Klein, Dana Blumenthal, Alan Knapp
37. Sierra Foothills REC (sier.us) - Eric Seabloom, Elizabeth Borer, W. Harpole
38. Smith Prairie (smith.us) - Jonathan Bakker, Janneke Hille Ris Lambers
39. Spindletop (spin.us) - Rebecca McCulley, Jim Nelson
40. Temple (temple.us) - Philip Fay, Jason Martina
41. Trelease (trel.us) - Andrew Leakey
42. Ukulinga (ukul.za) - Kevin Kirkman, Michelle Tedder

43. Val Mustair (valm.ch) - Anita Risch, Martin Schuetz
44. Yarramundi (yarra.au) - Sally Power, Raul Ochoa Hueso
